# Freeing P300-Based Brain-Computer Interfaces from Daily Calibration by Extracting Daily Common ERPs

**DOI:** 10.1101/2024.03.02.581675

**Authors:** Dojin Heo, Sung-Phil Kim

## Abstract

When people use brain-computer interfaces (BCIs) based on event-related potentials (ERPs) over different days, they often need to repeatedly calibrate BCIs every day using ERPs acquired on the same day. This cumbersome recalibration procedure would make it difficult to use BCIs on a daily basis. We aim to address the daily calibration issue by examining across-day variation of the BCI performance and proposing a method to avoid daily calibration. To this end, we implemented a P300-based BCI system designed to control a home appliance over five days in nineteen healthy subjects. We first examined how the BCI performance varied across days with or without daily calibration. On each day, P300-based BCIs were tested using calibration-based and calibration-free decoders (CB and CF), with a CB or a CF decoder being built on the training data on each day or those on the first day, respectively. Using the CF decoder resulted in lower BCI performance on subsequent days compared to the CB decoder. Then, we developed a method to extract daily common ERP patterns from observed ERP signals using the sparse dictionary learning algorithm. We applied this method to the CF decoder and retested the BCI performance over days. Using the proposed method improved the CF decoder performance on subsequent days; the performance was closer to the level of the CB decoder, with improvement of accuracy by 2.28%, 1.93%, 1.75%, and 3.86 % on the subsequent four days, respectively, compared to the original CF decoder. The method proposed by our study may provide a novel approach to addressing the daily-calibration issue for P300-based BCIs, which is essential to implementing BCIs into daily life.

## 1. Introduction

A brain-computer interface (BCI) enables people to directly interact with the external world by reading users’ intention from brain signals [1]. Brain signals for BCIs can be acquired non-invasively, making the BCI system more convenient to use [2]. According to strategies for inducing brain signal patterns that effectively represent users’ intention, BCIs can be categorized as active, passive, and reactive BCIs [3]. Among them, a reactive BCI draws on the brain responses to external stimuli, each being associated with a specific intention of users. The present study focuses on reactive BCIs using electroencephalography (EEG), and specifically those based on event-related potentials (ERPs). One of the most popular ERP components used for BCIs is the P300 component [4], which is generated by the oddball paradigm of stimulus presentation [5]. The P300-based BCI system has been extensively developed so far and applied to assist the living of clinical populations such as patients with amyotrophic lateral sclerosis (ALS) [6-8]. Moreover, it has recently been developed for non-clinical individuals with applications such as deception detection [9], playing games [10], and controlling home appliances [11], to name a few.

Yet, many challenges have still remained to be overcome before using the P300-based BCI system in daily life [12]. BCI-illiterate users, for instance, exhibit problems using P300-based BCIs to deliver their intention because their P300 signals are relatively faint [13]. BCI systems using dry electrodes are preferred to those using wet electrodes in real life due to their high usability, but such systems still confront the problem of low signal quality [14]. It is also important to tackle other issues such as distraction by multitasking [15, 16] and artifacts by natural movements [17]. Moreover, one of the most practical issues is time-consuming calibration necessary to set up P300-based BCIs for everyday use [18]. Accordingly, the users should undergo a laborious calibration before using the BCI system every day.

The necessity of daily calibration largely originates from the non-stationarity of EEG signals. It can be attributed to spatiotemporal neural dynamics linked to time-varying functional communications between brain regions [19]. In this regard, Nishimoto *et al*. demonstrated that EEG features extracted on two different days were distributed differently, representing the existence of intra-subject variability over days [20]. Similarly, Changolusia *et al*. reported spatiotemporal differences in ERPs across two different days. [21].

In addition to intrinsic spatiotemporal neural dynamics, other causes of the non-stationarity of P300 have been suggested. Users’ physiological conditions might fluctuate over time while using the P300-based BCI system. Physiological fluctuations arising from circadian rhythm, food consumption, physiological arousal, and fatigue could lead to variations in the P300 component [22]. Also, the habituation effect of repeatedly presenting stimuli could alter attentional states, resulting in a decrease in the amplitude of the P300 component [23-26].

Such variations in P300 (or more generally ERP) features on different days inevitably necessitate the daily calibration of P300-based BCIs where BCI systems are rebuilt everyday using EEG data obtained on the same day [27]. From the user’s perspective, daily calibration can be cumbersome, lowering self-motivation and making users feel uncomfortable using the P300-based BCI system.

To address the daily calibration issue, which is common in not only P300-based BCIs but also a variety of non-invasive and invasive BCIs, a few studies have proposed potential solutions. A study by Barachant and Congedo proposed a Riemannian manifold method to reduce the time for daily calibration of P300-based BCIs over six days [28]. Krauledat *et al.* investigated the long-term use of motor imagery based BCIs (MI-BCIs) by incorporating common spatial pattern (CSP) filters derived from a pre-existing signal set to account for daily variability [29]. Christensen *et al*. reported reduced error in classifying cognitive load levels from EEG when subjects performed multitasking over five days by adding only short-time EEG signals on each new day to recalibrate a decoder [30]. A study for EEG-based emotion recognition separated task-related EEG components being consistent over five days from background EEG components varying over days by using the robust principal component analysis (RPCA), and used the task-related components to enhance cross-day classification [31]. However, all of these studies did not avoid daily calibration completely. In contrast, other invasive BCI studies have proposed subspace methods to find low-dimensional manifolds of neuronal population activities that were relatively stable over days and used those manifolds to improve BCI control without calibration [32, 33].

These previous studies indicate that identifying and separating daily common components from daily varying ones could be effective in addressing day-to-day variability of EEG. Accordingly, we aim to address the daily calibration issue by taking an unsupervised approach to extract daily common components of ERP waveforms and to use them to build a calibration-free P300-based BCI system. Specifically, we propose using the sparse dictionary learning (SDL) method to extract daily common components. SDL, an algorithm to learn sparse representations of data, can describe observed ERPs as the linear combination of a set of template waveforms with sparse coefficients. Each template waveform, called an “atom”, represents a common waveform embedded across all the observations. More than one atom may be necessary to represent observed ERPs and a collection of atoms is called a “dictionary”. The sparse coefficients enables the representation of ERPs only by significant atoms [34].

SDL has been recently used for EEG studies, including the detection of EEG patterns related to epilepsy [35] or schizophrenia [36], and feature extraction in MI-BCIs [37]. But the primary purpose of using SDL in these studies was to enhance classification performance. On the other hand, Morioka *et al*. employed SDL to resolve the inter-subject variability in MI-based BCIs by extracting spatially common signal patterns across all subjects. [38]. Nishimoto *et al.* also employed SDL to extract personal EEG features that were consistent across different times of the day, specifically between morning and afternoon sessions [20].

For ERPs, we can anticipate that a set of atoms determined by SDL would sufficiently represent ERP waveforms consistent over days while suppressing daily varying components [39, 40]. However, we may need across-day ERPs to learn daily common atoms. This is still problematic because we should collect data over different days via a calibration procedure. To circumvent this problem, we aim to find atoms only using ERPs on the first day and utilize them to completely avoid daily calibration for the use of P300-based BCIs on the following days.

The specific aims of this study are two folds. First, this study aims to assess whether the P300-based BCI performance deteriorates over days if daily calibration is not conducted. Second, this study aims to propose a method based on SDL to improve the consistency of P300-based BCI performance over days without daily calibration. We use SDL to acquire a dictionary that encompasses a set of atoms capable of capturing ERP waveforms robust to fluctuations over time. This dictionary is learned from the data collected on the first day. As such, we expect that the reconstructed ERPs on the following days, utilizing the dictionary learned on the first day and their respective coefficients determined on each day, would exhibit ERP waveforms similar to those observed on the first day.

## 2. Experiment

### 2.1 Subject

Nineteen healthy subjects (11 male, ages 19-27 years old with a mean of 23.47±2.61) participated in the experiment. Except for one subject, no one had experienced using the P300-based BCI system prior to the experiment. No subject was taking medication for neurological or mental illness. All subjects had normal or were corrected to normal vision. To minimize potential nuisance factors, subjects were instructed to limit caffeine intake to no more than 300mg on the day of the experiment and to avoid excessive alcohol consumption the day before. All subjects also had a minimum of 4.5 hours of sleep before participating in the experiment.

### 2.2 Experimental Protocol

Subjects participated in the experiment for a total of five days. The experiment was conducted for the first four days in a row, then for another day after a break. This break, which lasted nine to eleven days, was designed to examine if a habitual response could be relieved by taking days off (Figure1(A)). Subjects were guided not to participate in any other BCI experiments during the break. The experiment always began at the same time of the day for each subject.

### 2.3 Stimuli

On a black background, four light blue box-shaped icons were continuously displayed in each of the four corners of the computer screen. Each icon expressed simple shapes (i.e., square, triangle, hexagon, and star) in the calibration session or shapes related to control commands (i.e., on, off, color change, or brightness change) of an electronic lighting device (ELD, Phillips Hue 2.0, Phillips, Inc., The Netherlands) operated by the P300-based BCI system in the test session (see Section 2.4 below). Each icon served as a stimulus by turning its color to yellow (i.e., highlighted) (Figure 1(B)). The size of each icon was 5.5°×6.0°. The size of computer screen was 42.2° × 24.5°. The distance from the computer screen to subjects was maintained at 70 cm.

**Figure 1.**
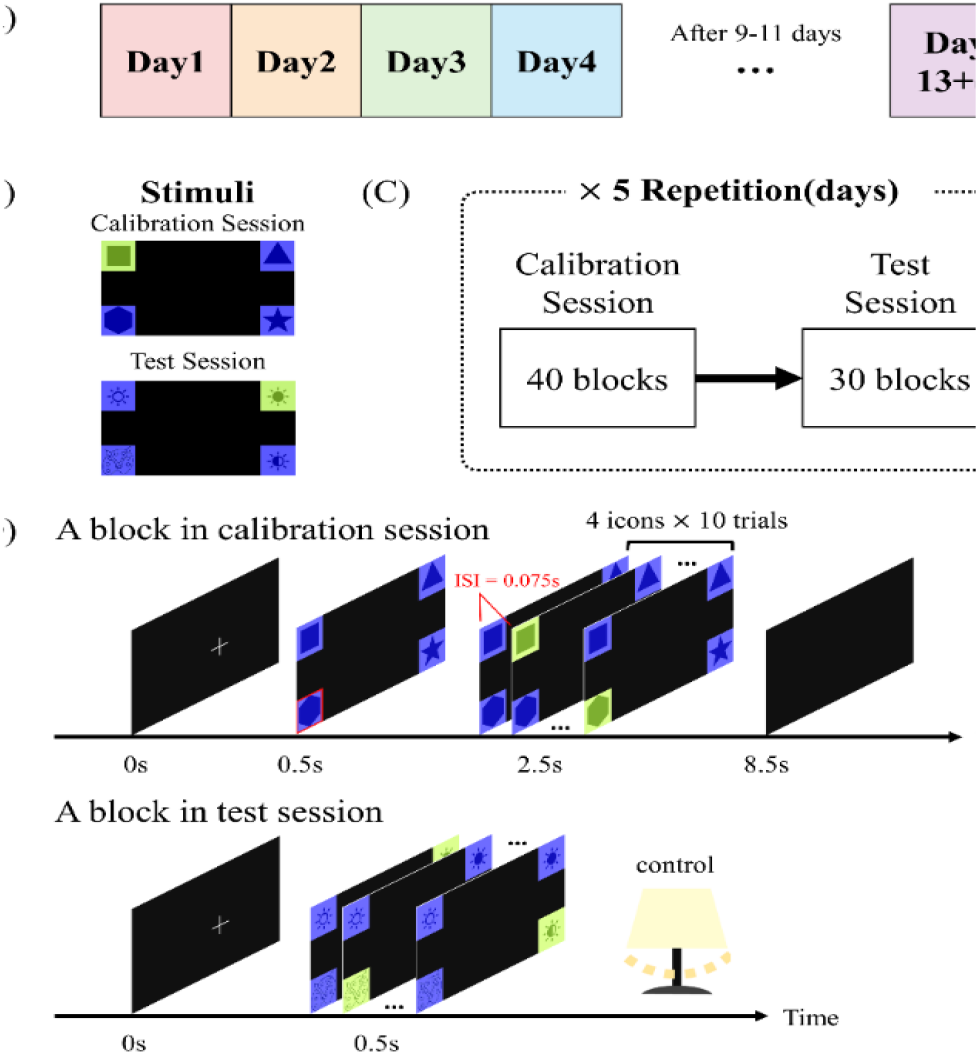
(**A**) An overall experimental protocol of this study on daily calibration for brain-computer interfaces (BCIs). The experiment was carried out over a period of four consecutive days, followed by an additional day after a break lasting nine to eleven days. Each day was marked by different colors. (**B**) Illustrative stimuli designed for calibration and test sessions. In the calibration session, the stimuli consisted of basic icons such as squares, triangles, hexagons, and stars, whereas in the test session, each icon was associated with the control command of an electronic lighting device. In both sessions, a stimulus was presented by turning the corresponding icon from blue to light green. (**C**) On each day, subjects performed the calibration session including 40 blocks followed by the test session including 30 blocks. (**D**) A block consisted of a sequence of stimuli presented to subjects. During the calibration session, a block began with a white fixation cross for 0.5s, followed by the notification of a target stimulus to which subjects paid attention for 2s, marked by a red contour square. Subsequently, each of four stimuli was pseudorandomly presented for 0.075s with an inter-stimuli interval (ISI) of 0.075s. Every stimulus was presented 10 times. Afterward, a black screen was displayed for 1s. A block in the test session was the same as that in the calibration session except that the step of presenting a target stimulus was omitted. Instead, the experimenter verbally informed subjects of a target stimulus before each block. A block in the test session ended with an additional step wherein the electronic lighting device operated a function selected by the BCI output.

### 2.4 Task

Subjects performed the visual oddball task to build and run the P300-based BCI system. On each day of experiment, subjects performed a calibration session to build the system and a test session to run the system online. The calibration session included forty blocks and the test session included thirty blocks of the oddball task. The test session always followed the calibration session (Figure 1(C)).

In the calibration session, a block began with the fixation period in which a white cross was shown at the center of the screen for 0.5s. The size of the fixation cross was 2.0° × 2.0°. Then, the four squared icons appeared at each corner with the disappearance of the fixation cross. At the same time, a red squared contour appeared on top of one of the icons for 2s to inform subjects of a target stimulus to which subjects should pay attention. Each icon was marked in red across blocks pseudorandomly, ensuring that each icon was selected uniformly as a target. Then, each icon was highlighted for 0.075s sequentially in a random order with 0.075s interstimulus interval (ISI). As such, one round of highlighting the four stimuli took 0.6s. This design of stimulus presentation time (0.075s) and ISI (0.075s) was determined as the minimum amount of time needed to present visual stimuli in the P300-based BCI system by previous work [41]. We repeated the round of stimulus presentation ten times with no break between rounds, which took 6s. Accordingly, there were ten trials of target stimulus presentation and thirty trials of non-target stimulus presentation per block (the ratio of the number of target stimuli to that of non-target stimuli was 1:3). In each block, subjects were instructed to count the number of times the target stimulus was presented, which was intended to help them sustain attention. Finally, the block ended with a black screen for 1s (Figure 1(D), top). The P300-based BCI system was built using the data acquired from the calibration session.

In the test session, a block was the same as in the calibration session, except that the target was indicated not by the red contour but by a verbal instruction from the experimenter before starting the block. The icons were also changed to control command shapes. The instruction was predetermined before the test session to ascertain that subjects attempted to control each of the four ELD functions in a random order without consecutively repeating the same control. After giving the instruction, the experimenter ran a block from fixation to the ten rounds of stimulus presentation (Figure 1(D), bottom). Then, the P300-based BCI system was operated to generate one of the four control commands for the ELD. The command was directly sent to the ELD to control the device in real time. The control result was immediately fed back to subjects.

## 3. Signal acquisition and classification

### 3.1 EEG signal acquisition

EEG signals were acquired from 31 active wet electrodes (Fp1, Fpz, Fp2, F7, F3, Fz, F4, F8, FT9, FC5, FC1, FC2, FC6, FT10, T7, C3, Cz, C4, T8, CP5, CP1, CP2, CP6, P7, P3, Pz, P4, P8, O1, Oz, and O2) attached to the scalp based on a standard head-mounted EEG cap following the 10-20 system of American Clinical Neurophysiology Society Guideline 2. Electrodes for ground and reference were positioned at the mastoids of the left and right ears, respectively. Impedance for all electrodes was kept below 10 kΩ throughout the EEG measurement. The EEG signals were amplified by a commercially available EEG amplifier (actiCHamp, Brain Product GmbH, Gilching, Germany) and sampled at 500 Hz.

### 3.2 EEG signal preprocessing and ERP extraction

To remove artifacts, the measured EEG signals were preprocessed via the following steps: 1) High-pass filtering by a 4^th^-order Butterworth filter above 0.5 Hz was applied to the signal. 2) Electrodes that contained EEG signals showing correlations lower than 0.4 with EEG signals at more than 70% of all other electrodes were rejected [42]. As a result, the average number of 2.84±1.91 electrodes were rejected, with a maximum of 11 electrodes, across subjects. 3) EEG signals of the remaining electrodes were re-referenced by common average reference (CAR). 4) EEG signals were low-pass filtered by a 4th-order Butterworth filter below 50 Hz. 5) The artifact subspace reconstruction (ASR) method was applied to remove artifacts by setting the cutoff parameter as 10 [43, 44]. 6) Finally, EEG signals were low-pass filtered again below 12 Hz by a 4th-order Butterworth filter.

To obtain ERPs, the preprocessed signals were epoched by a time window of -0.2 to 0.6s after stimulus onset. As there were 10 presentations of each stimulus per block, 10 epoched EEG signals were averaged for each stimulus, thereby producing 4 ERPs from each block. Baseline correction was applied to these ERPs by subtracting the mean and dividing by the standard deviation of the baseline amplitudes where a baseline period was determined to be -0.2 to 0s. As such, we obtained a total of 160 ERPs, 40 for target and 120 for non-target stimuli, at each electrode during the calibration session and 120 ERPs (30 targets and 90 non-targets) during the test session.

### 3.3 Decoding

A decoding algorithm in the P300-based BCI system was designed to classify given ERP data into a target or non-target class (binary classification). ERP data for two classes obtained from the calibration session were used to train a classifier, where the Support Vector Machine (SVM) with a linear kernel was used in this study. Let a matrix 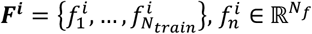, denote a feature set composed of training ERP data collected on day *i, i* = 1, 2, …, 5. Here, days 1 to 4 refer to the first four consecutive days and day 5 refers to the last day after a break (see above). *N*_*f*_ denotes the dimensionality of a feature vector, *f*^*i*^, where *n* = 1, …, *N*_*train*_ indexes a training sample and *N*_*train*_ is the number of training samples (*Ntrain* = 160 in this study). The feature vector was created by concatenating the ERPs across all the electrodes. We sampled baseline-corrected ERP amplitude values at intervals ranging from 0.15 to 0.60s. Thus, the number of ERP values (denoted as *N*_*t*_) at each electrode was 225 ( 0.45*s* × 500 *Hz*). The number of electrodes (denoted as *N*_*e*_) was not fixed since the number of rejected electrodes varied across subjects and days. The dimensionality of the feature vector was determined as *N*_*f*_ = *N*_*t*_ × *N*_*e*_. For instance, if no electrode was rejected, *N*_*f*_ = 225 × 31 = 6,975. The class label in SVM was set as 1 for target and -1 for non-target. Given the feature set ***F*^*i*^** for training, the linear SVM classifier was trained to discriminate between target and non-target stimuli. The penalty parameter in SVM was set as ***C*** = 1.

After training, the normal vector to the hyperplane, *ω*^*i*^ in the linear SVM built on the feature set on day *i* was used to classify each of *N*_*test*_ feature vectors in the test session on day *j* into the target or non-target class (*N*_*test*_ is the number of test samples and equal to 120 in this study). Specifically, after each block of the test session, we predicted the target stimulus (*s*\*) out of four based on the SVM outputs by:

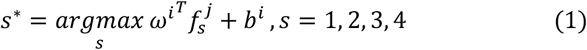

where *s* indicates each of the four stimuli, 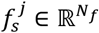 is the feature vector on day *j* corresponding to a stimulus *s*, and *b*^*i*^ is a bias term. The stimulus *s*\* was decoded as target and other three stimuli as non-target in each block.

### 3.4 Test of BCIs with and without calibration

In this study, we constructed two different decoding models depending on daily calibration. In one decoding model, the SVM classifier trained using data from the calibration session on day *i* was used to decode data in the test session on the same day (i.e., *j* = *i*) – termed a calibration-based (CB) decoder. In the other model, the SVM classifier trained using data from the calibration session on day 1 was used to decode data in the test session of day *j* (*j* = 1, 2, …, 5) such that no calibration was necessary except for day 1 – termed a calibration-free (CF) decoder (Figure 2). In this case, *j* ≠ *i* except that *j* = 1.

**Figure 2.**
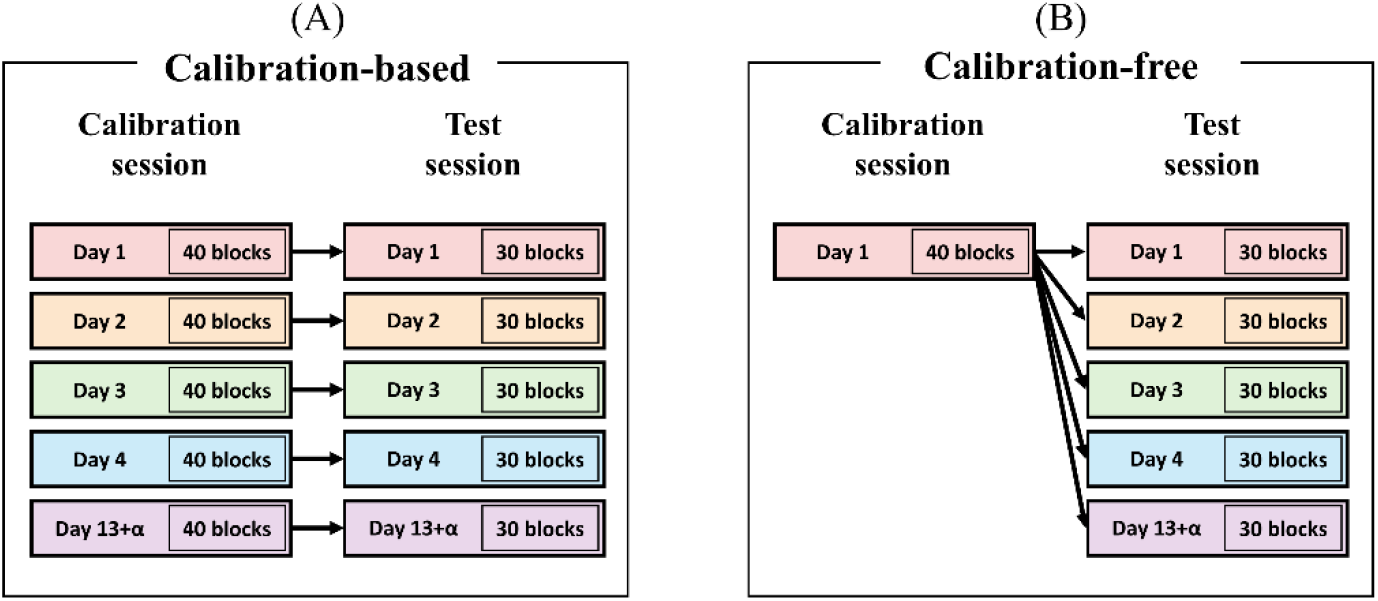
Design of two decoding models depending on calibration. (**A**) In the calibration-based (CB) decoding model, a decoder was trained using EEG data from the calibration session and used for the test session on the same day. Black arrows indicate association of training and testing data for the decoder. (**B**) In the calibration-free (CF) decoding model, a decoder was trained using EEG data from the calibration session on day 1 and used for the test sessions on each day afterward.

Note that the CB decoder was tested online whereas the CF decoder was tested offline, as we opted for the CB decoder in our online BCI experiments to acquire daily calibration data for further analyses. In a different approach, we could have tested the CF decoder first followed by the calibration session and the CB decoder on days 2 to 5. Yet, in this case, we were concerned with a possible learning effect of the initial CF test on the subsequent CB test. So, we opted for applying the CB decoding only on each day and analyzing the CF decoding offline to analytically compare two decoders.

Given that the rejected electrodes could differ daily in each subject, it was likely that *N*_*f*_ would not be equal between the calibration and the test sessions in the CF decoding. To address this, we included only those electrodes that were not rejected during the preprocessing procedures of day 1 nor day *j* to test the CF decoder on day *j*.

## 4. Calibration-free solution

In this section, we first describe conventional sparse dictionary learning (SDL) and then introduce our proposed method that modifies SDL for our purpose.

### 4.1 Sparse dictionary learning

Consider a set of ERP waveforms 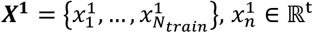, at a given electrode collected from the calibration session on day 1. From ***X*^1^,** SDL learns a dictionary 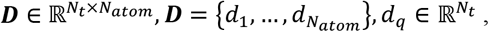, where *N*_*atom*_ is the number of atoms, and a set of coefficient vectors 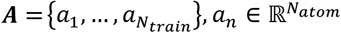 such that the linear combination of ***D*** by ***A*** reconstructs ***X*^1^** as accurately as possible [45]. SDL can be formulated as the following optimization problem:

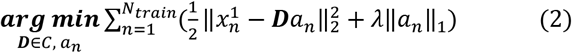

where *λ >* 0 controls a balance between the sparsity of *a* and the minimization of reconstruction error, which is set to 0.8 through a grid search process, where the best averaged accuracy for subjects was identified within the range of 0.1 to 1 at intervals of 0.1. This value aligns with the recommended range as indicated in prior research study [46]. ‖·‖_*p*_ denotes *l*_*p*_-norm and ***C*** is a constraint for ***D*** preventing the atoms from having arbitrarily large values such as:

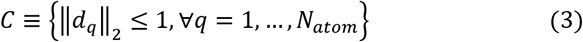

To optimize ***D*** and ***A*** together, SDL repeatedly minimizes the objective function by adjusting one while fixing the other and vice versa until convergence. In each repetition, each ***D*** and ***A*** optimizations are performed by block coordinate descent [47] and least angle regression (LARS) [48], respectively. *N*_*atom*_ is a hyperparameter that needs to be pre-determined before learning. Here, we employed the “mexTrainDL” and “mexLasso” functions from the SPAMS toolbox using the MATLAB program that implemented online dictionary learning algorithm developed by Mairal *et al.* [45].

### 4.2 Residual sparse dictionary learning

We assumed that a set of “ERP atoms” in the dictionary would be more robust to variation of observed ERPs. Thus, using reconstructed ERPs from sparsely combining ERP atoms would maintain essential ERP waveforms. However, using the conventional SDL for P300-based BCIs faces two challenges. First, it is unknown a priori how many atoms are necessary in each subject. The number of atoms is usually determined heuristically or by grid searches. However, this approach may not be practical for online BCI systems. Previous studies have recognized the issue of an unknown dictionary size and attempted to address this issue [49-52]. Second, certain atoms can be redundant as the conventional SDL places no constraints on minimizing redundancy between atoms. Minimum redundancy between atoms is critical to a dictionary for ERPs because a small change in the latency of ERPs over multiple EEG electrodes could generate different atoms, which may increase a chance to miss other important atoms.

Therefore, we developed a modified version of SDL in this study, which is called residual sparse dictionary learning (rSDL). A basic idea of rSDL is that each atom is constructed one by one from residuals of the estimation of observed data ( ***X*^1^**) by preceding atoms (***D***) and corresponding sparse coefficients (***A***). This approach is similar to other learning methods such as cascade neural networks [53], deflation methods [54], and orthogonal matching pursuit (OMP) [55]. Learning stops when remaining residuals meet stopping criteria (see below for details). In this way, we can automatically determine *N*_*atom*_ during learning, with no need to predetermine it a priori. Also, the resulting atoms can be more independent to each other as we learn each atom using residuals, similar to commonly known Gram-Schmidt orthogonalization process [56]. Therefore, rSDL may circumvent the challenges in SDL and yield atoms more appropriate to find essential ERP patterns.

An outline of rSDL is described in Figure 3(A) and the detail procedure of rSDL is illustrated in algorithm 1. Given 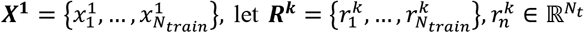 be a set of residuals at the *k*-th iteration of rSDL. In the beginning of the iteration with *k* = 1, we initialize ***R*^0^** = ***X*^1^**. Then, we infer *d*_*k*_ and 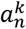 for *n* = 1, …, *N*, by solving the following optimization problem with 1,000 times of loop (T=1000),

**Figure 3.**
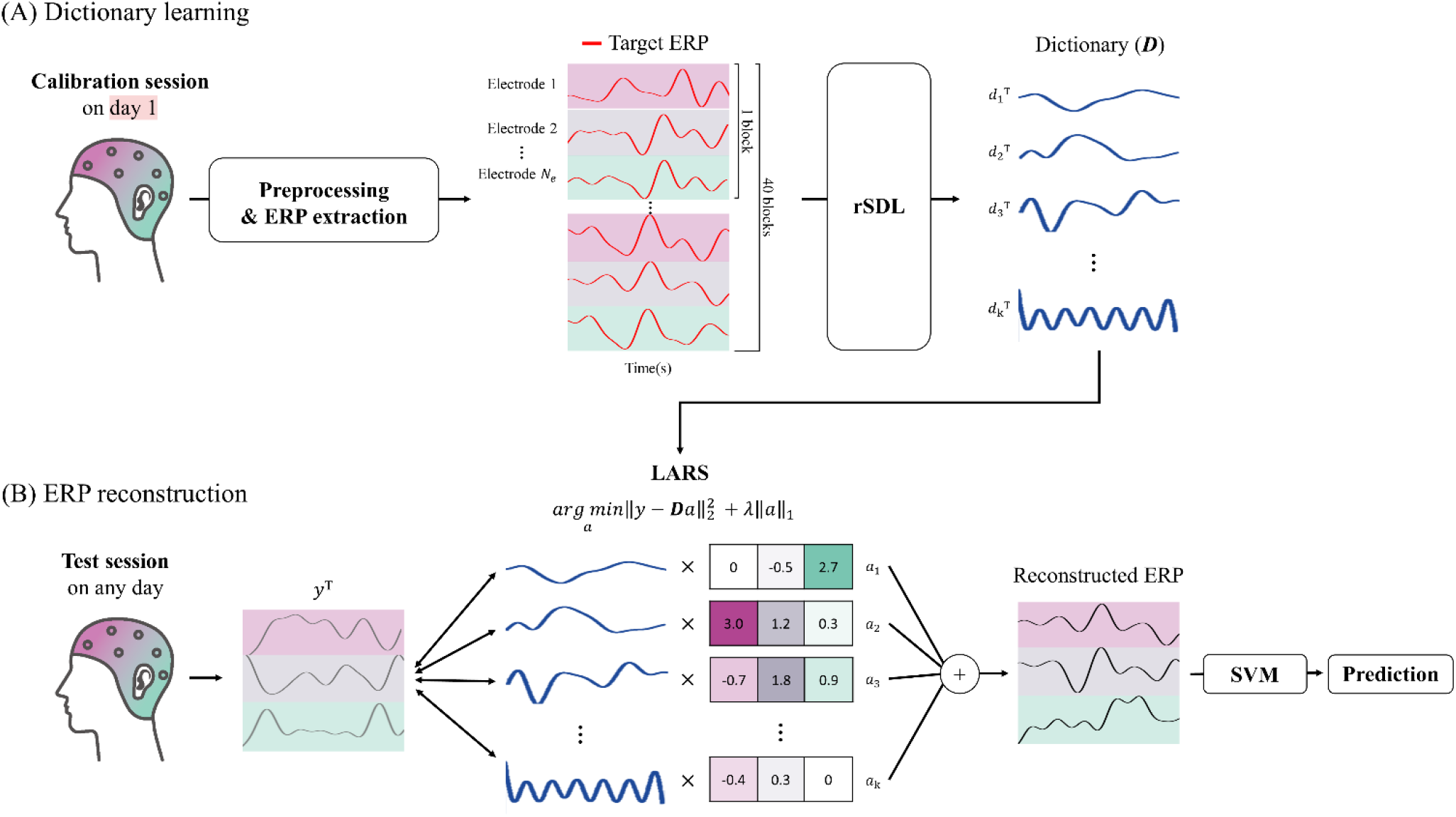
Outline of the proposed model for calibration-free decoding over days. The proposed model was applied on an individual level. (**A**) Illustration of dictionary learning. The EEG signals acquired during the calibration session on day 1 underwent preprocessing and extraction of ERPs in response to target stimuli (target ERPs). The target ERPs extracted at each of *N*_*e*_ electrodes from 40 blocks were utilized to learn a dictionary using the residual sparse dictionary learning (rSDL) method. Using the rSDL method, we learned *k* atoms (*d*_*i*_, *i* = 1, …, *k*) each representing common ERP patterns. These k atoms constituted a dictionary *D*. (**B**) Illustration of the application of the learned dictionary to calibration-free decoding. During the test session on any day, *D* was used to derive coefficients (*a*) using the least-angle regression (LARS) algorithm for given ERPs (*y*) for all electrodes. *D* and *a* were then linearly combined to reconstruct ERPs. The reconstructed ERPs were utilized to predict a target stimulus via support vector machine (SVM) that was trained with the calibration session data on day 1.

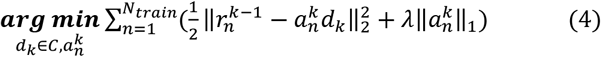

using SDL with *N*_*atom*_ = 1, as described in Section 4.1 above. Since there is a single atom, 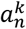 is a scalar. After inferring *d*_*k*_ and 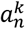, we update the residuals as 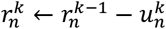, where 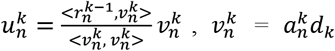, to impose orthogonality between 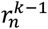 and 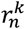. Then, the next iteration is repeated by increasing *k* to estimate *d*_*k*_ and 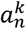 until a pre-set criterion for convergence is met. Note that *d*_*k*_ and 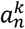 are estimated independently of {*d*_1_, …, *d*_*k*−1_} and 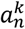. After the end of iterations, we concatenate *d*_*k*_ and 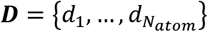 from every iteration to construct 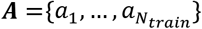and 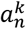, where *a*^*k*^ from the *k*-th iteration constitute the *k*-th row of ***A***.

A stopping criterion in this study was set as *l*(*k*) − *l*(*k* − 1) < 10^−4^, where *l*(*k*) was a loss function calculated at the *k*-th iteration and defined as the ratio of the root sum square of 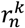 to the root sum square of 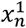 over the index *n*. In this study, 13.71±0.47 atoms were learned on average across subjects.

#### Algorithm 1. Residual-based sparse dictionary learning (rSDL)

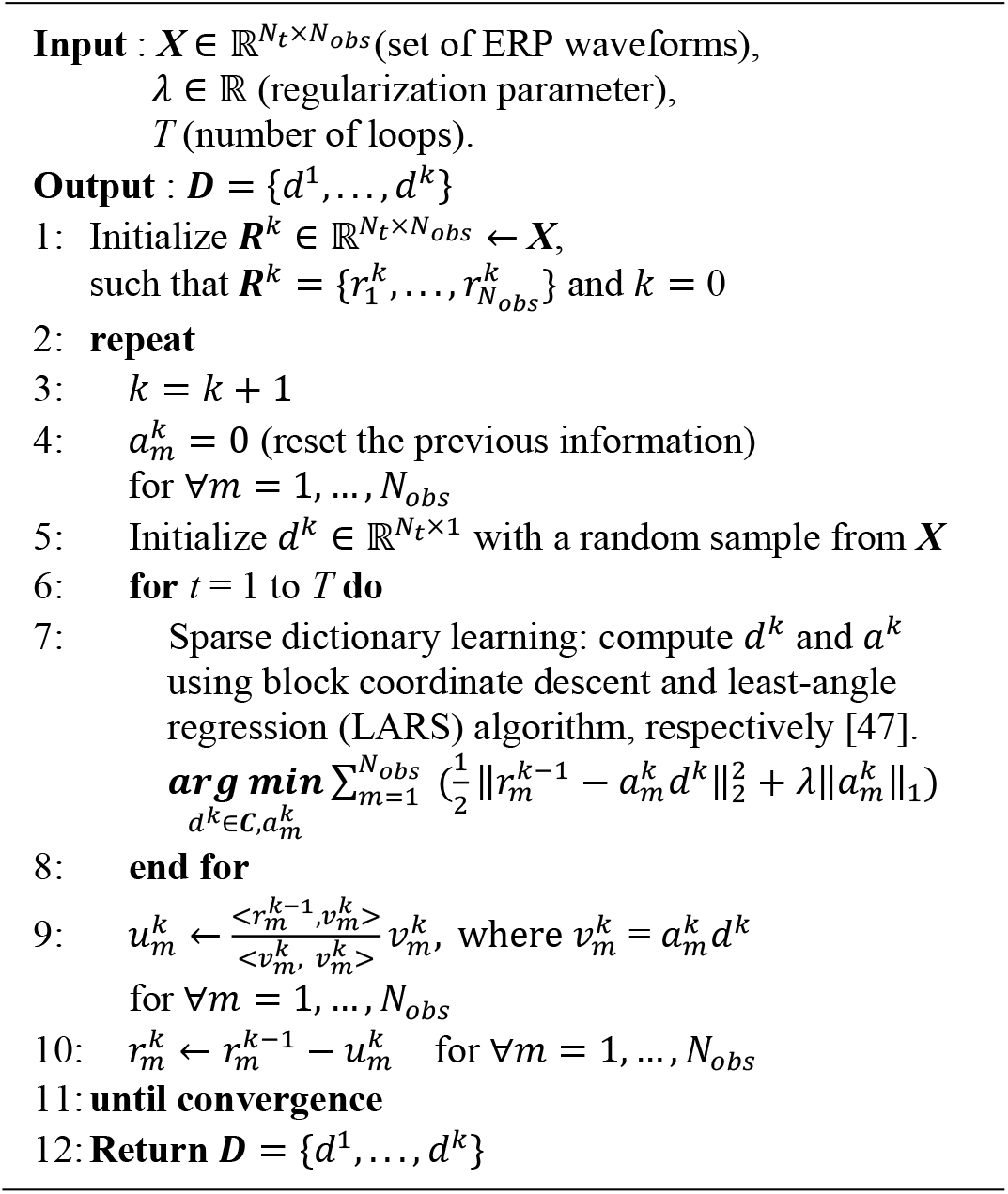

We estimated ***D*** using ERPs in response to target stimuli only fas we intended to reconstruct ERP waveforms representing target stimuli using the dictionary. As ERPs in response to non-target stimuli often exhibit random oscillations in the signal, the learned dictionary from ERPs in response to non-target stimuli may inadvertently incorporate atoms containing random oscillations. We attempted to minimize the acquisition of such atoms, as their presence could potentially decrease the consistency of reconstructed ERPs across days.

Then, we reconstructed ERPs by multiplying the learned atoms by corresponding coefficients. These reconstructed ERPs, concatenated across all electrodes, were used to train the linear SVM in the CF decoder. For the test data, we estimated the coefficient vector *a* for each test sample using ***D***. Let 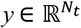 denote a new test ERP sample. Then, we used LARS to solve the following problem to estimate *a*:

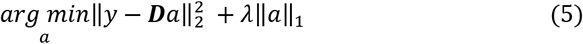

In this way, the coefficient for each test sample in the test session on any day (i.e., day 1, 2, 3, 4, and 13+α) was estimated. Using the estimated coefficients, we reconstructed ERPs, decoded using the linear SVM and predicted a target stimulus in a given test block (Figure 3(B)).

## 5. Data analysis

### 5.1 BCI performance evaluation

We assessed the performance of a decoding model using accuracy, which was defined as the percentage of correct test blocks among the whole test blocks. Hereafter, we denote different decoding models by {decoder type}:{ERP type}; the decoder type is either CB or CF, and the ERP type is either original ERPs (org) or reconstructed ERP by rSDL (rSDL). Essentially, we analysed three decoding models: CB:org, CF:org and CF:rSDL in this study.

First, we compared the performance between CB:org and CF:org to examine how the absence of daily calibration influenced decoding performance. Note that performance was identical for CB:org and CF:org on day 1. Thus, we assessed the performance of CB:org and CF:org on subsequent days (2, 3, 4, 13+α) to examine the main effect of decoders on decoding accuracy (Figure 4).

**Figure 4.**
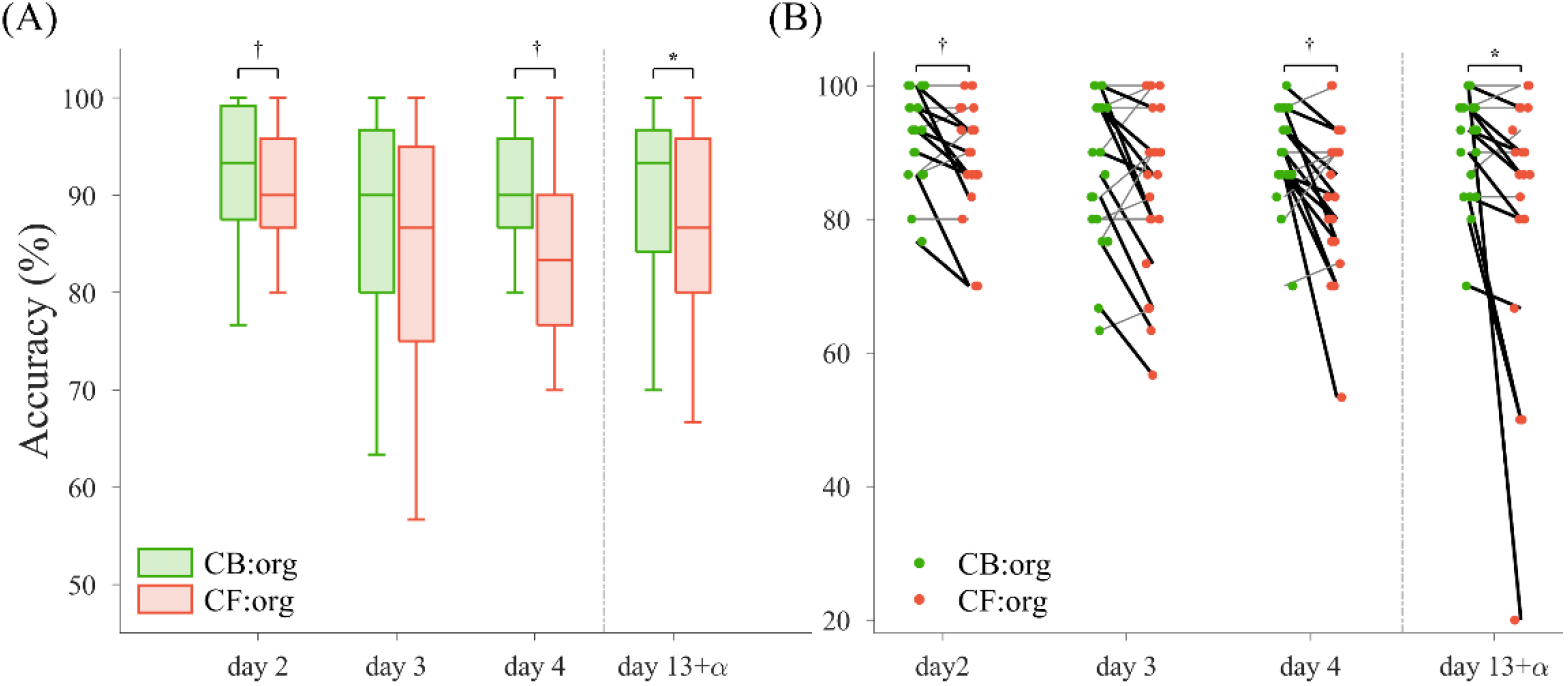
Performance of calibration-based (CB) and calibration-free (CF) decoding models based on original ERP features (original ERPs: org) without using the proposed method. (**A**) The box plot illustrates the median (represented by the horizontal line inside the box), as well as the 25th and 75th percentiles (depicted by the box) with a range of ± 1.5 times the interquartile range (shown as whiskers) of decoding accuracy over subjects on day 2 to day 13+α. Decoding accuracy was evaluated for both the CB (CB:org, green) and the CF (CF:org, red) decoders within subjects. (**B**) Paired line plots to compare the accuracy of two decoding models within subjects. Each dot on the plot represents the accuracy of a subject on a given day for CB:org (greed) or CF:org (red). Thick black lines indicate a decrease in accuracy from CB:org to CF:org, while thin gray lines indicate an increase or no change. Vertical gray dashed lines denote a break between the fourth and fifth days which ranged 7 to 9 days (with α of 0, 1, or 2 days). Statistical significance level was p<0.1 (†) or p<0.05 (*), as determined by a two-sided Wilcoxon signed-rank test with Holm-Bonferroni correction.

Next, we evaluated the effect of the proposed model by examining differences in decoding accuracy. We assumed that the proposed model would improve performance of the CF decoder closer to the CB decoder, leading to smaller differences in decoding accuracy. The difference in decoding accuracy was calculated by subtracting the accuracy of CF:org from that of CB:org as well as the accuracy of CF:rSDL from that of CB:org (Figure 5).

**Figure 5.**
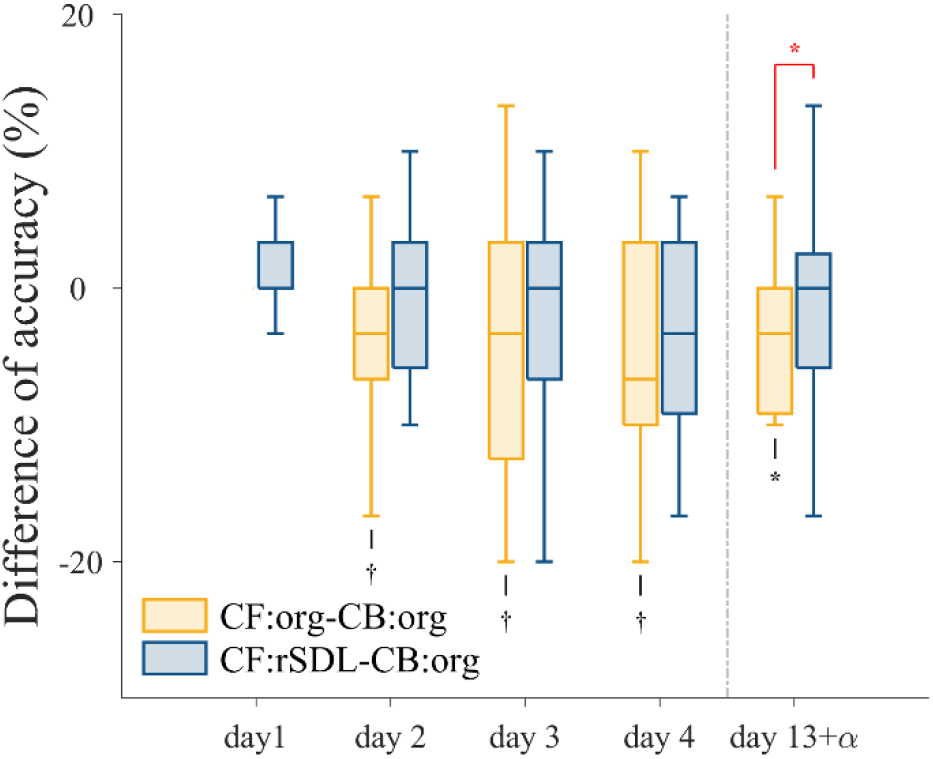
The difference in accuracy of the proposed calibration-free model (CF:rSDL) or the original calibration-free model (CF:org) relative to that of the calibration-based model (CB:org). The box plots illustrate the median (the horizontal line inside the box), as well as the 25th and 75th percentiles (range of the box) with a range of ± 1.5 times the interquartile range (whiskers) of decoding accuracy differences over subjects. The yellow and blue bars indicate the accuracy difference distributions for CF:org-CB:org and CF:rSDL-CB:org, respectively. Note that CF:org-CB:org is always 0 on day 1 as CF:org and CB:org used the identical training and testing data on day 1. A gray dashed line indicates a discontinuity between days (*α* = 0, 1, or 2 days). Significant differences between the proposed and original models are indicated by red asterisks (*), p<0.05. Black asterisks (*, p<0.05) and dagger (†, p<0.1) represent significant and marginally significant differences from zero for individual distributions (two-sided Wilcoxon signed-rank test with Holm-Bonferroni correction).

### 5.2 Feature evaluation

For the purpose of visual inspection, we visualized feature distributions using the t-distributed stochastic neighbour embedding (t-SNE) by lowering the dimensionality while preserving neighbouring structures. Feature vectors from the test sessions on all days were combined with those from the calibration session on day 1 to be used as input to t-SNE. Then, we extracted low-dimensional features using t-SNE for each ERP type (i.e. original and rSDL-reconstructed). We included the calibration session on day 1 here as the performance of the decoder was associated with distinction of feature distributions between the test and calibration sessions. After dimensionality reduction via t-SNE, we only collected the low-dimensional feature vectors in the test session on each day. As such, we created five low-dimensional feature distributions via t-SNE: test data on day 1 & calibration data on day 1, test data on day 2 & calibration data on day 1, …, test data on day 13+α & calibration data on day 1. We compared the feature distributions between original ERPs and reconstructed ERPs by rSDL (Figure 6).

**Figure 6.**
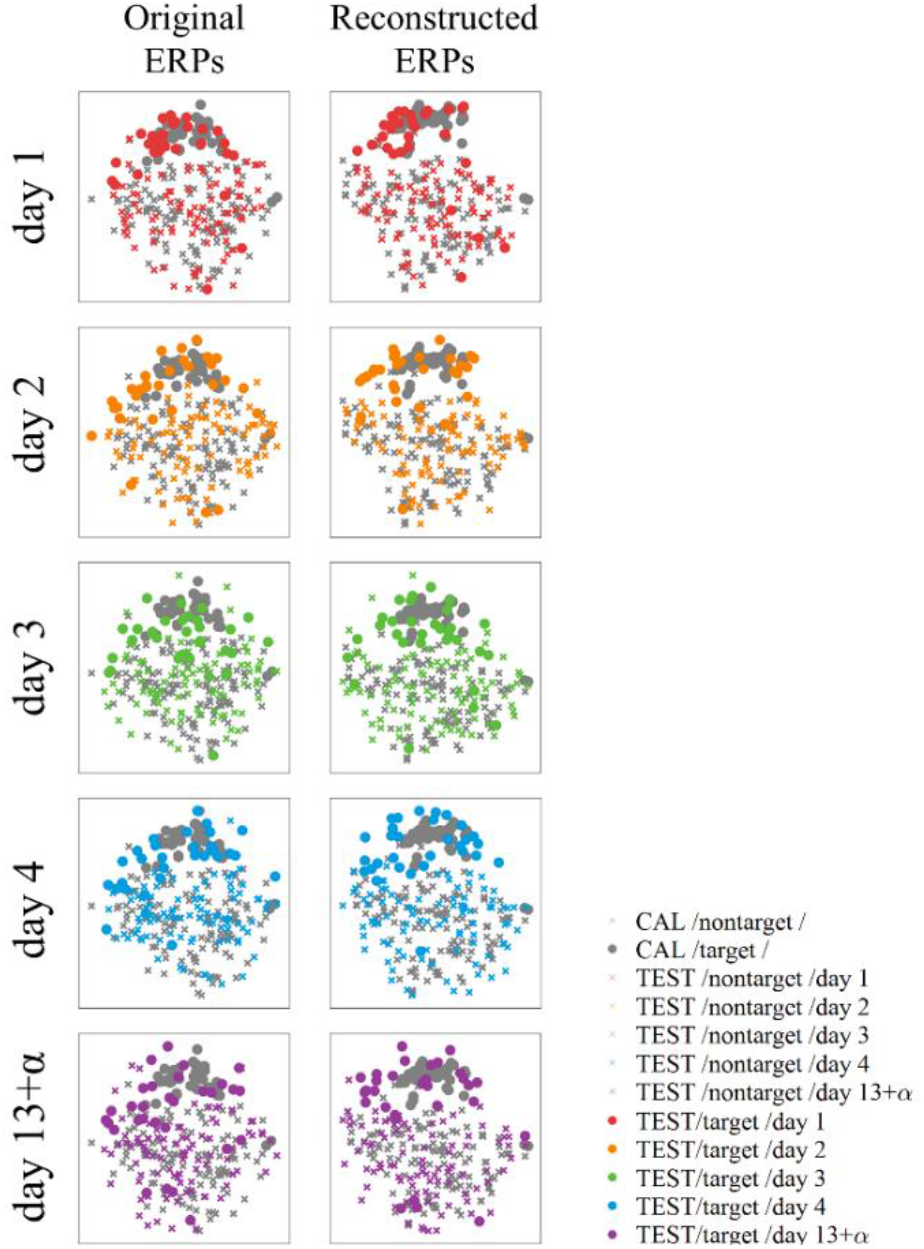
Example visualization of feature vectors on a low-dimensional space created by t-SNE. The feature vectors were extracted from either the original ERPs (left) or the reconstructed ERPs by rSDL (right) in a representative subject (subject 1). For each type of ERP, a decoder was trained using the feature vectors (gray) in the calibration session (CAL) on day 1 and tested on each day: day 1 (red), 2 (orange), 3 (green), 4 (blue), and 13+α (purple). Filled circles signify ERP features in response to a target and crosses does in response to a nontarget stimulus. If features distinguish target and nontarget perfectly, the clusters of circles and crosses would be completely separated from each other.

We examined the stability of ERP features across days by calculating the cosine distance of feature vectors between days [57]. A pair of feature vector sets, denoted as 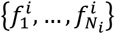 and 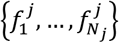, were considered, where the number of the samples (i.e. blocks) is equal to *N*_*i*_ for day *i* and *N*_*j*_ for day *j*, respectively. We equalized the dimensionality of the feature vectors from both days by using only those electrodes that were not excluded during the preprocessing of both day *i* and day *j*. Given a pair of feature vectors, the cosine similarity was calculated as follows:

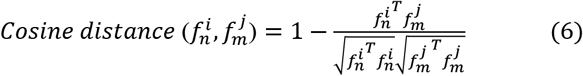

In our analysis, we calculated the cosine distance between a feature vector in response to target stimuli in the calibration session on day 1 and that in response to target stimuli in the test session of day *j*. As there were *N*_*i*_ = 40 feature vectors in the calibration session and *N*_*i*_ = 30 in the test session, we obtained 1,200 cosine distance calculations from all possible pairs. Then, we averaged them to obtain a final cosine distance between day 1 and day *j* for each subject (Figure 7).

**Figure 7.**
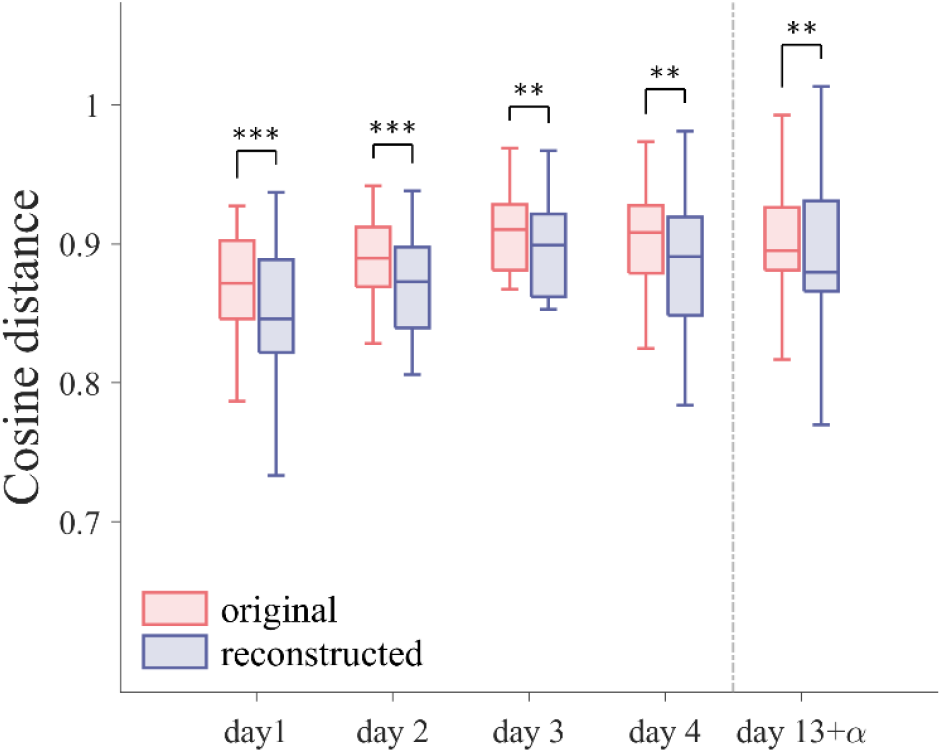
Comparison of the averaged cosine distance between the training and testing feature vectors using the original ERPs (red) or the reconstructed ERPs (blue). The cosine distance between feature vectors of the calibration session and those of the test session were calculated and averaged over all possible pairs (= 1,200) for each subject on each day. The box plots illustrate the median (the horizontal line inside the box), as well as the 25th and 75th percentiles (the range of the box) with a range of ± 1.5 times the interquartile range (whiskers) of the distribution of the averaged cosine distance over subjects. A gray dashed line indicates a discontinuity between days (*α* = 1, 2, or 3 days). Significant differences are denoted by asterisks (** p<0.01 and *** p<0.001, two-sided Wilcoxon signed-rank test with Holm-Bonferroni correction).

### 5.3 Signal-to-noise ratio analysis

We examined variation in the quality of ERPs across days in terms of the signal-to-noise ratio (SNR). A signal here was defined as the average of target ERPs over blocks in the calibration session (i.e., 40 blocks). We first obtained the average of target ERPs over blocks and then calculated the variance of averaged ERP over time to estimate the signal power. Afterward, we subtracted this averaged ERP from each of 40 target ERPs for each block to have residual ERPs. Then, we calculated the variance of individual residual ERPs over time for each block. The noise power was determined by the mean of the variance values over blocks [58]. SNR was estimated as a ratio of the signal power to the noise power for each electrode on each day (Figure 8).

**Figure 8.**
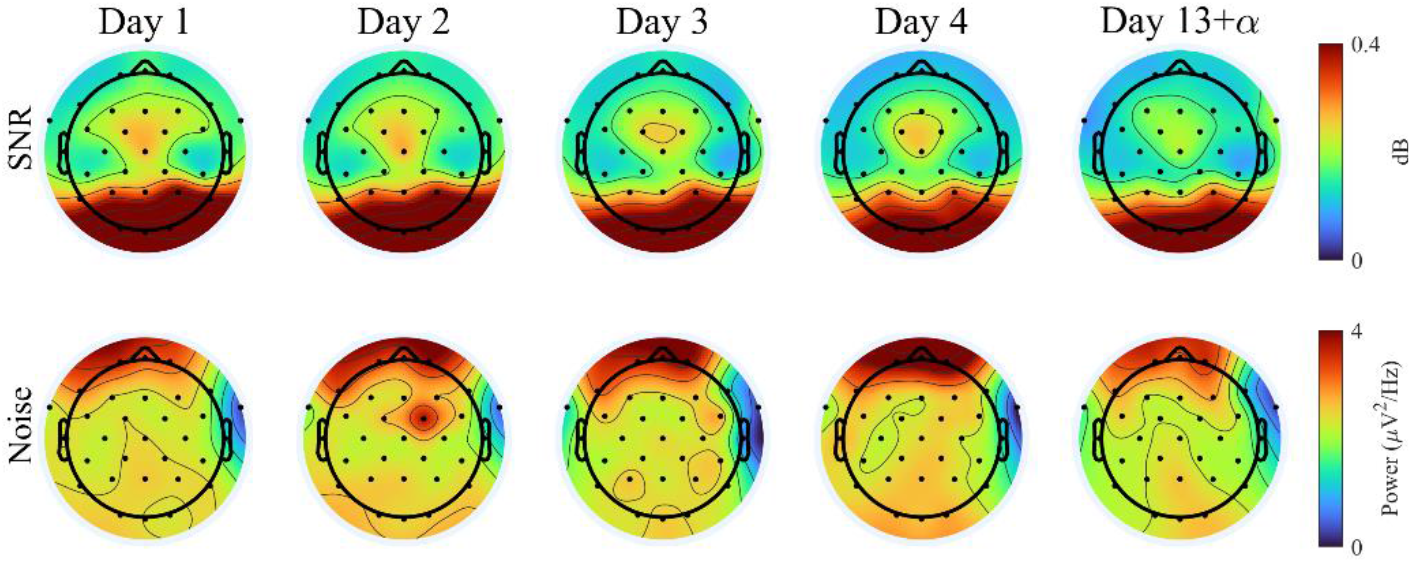
Topographies illustrate signal-to-noise ratio (SNR) (top) as well as noise power (bottom) of ERPs in response to target stimuli during the calibration session across days. Noise power was defined as the variance of residual ERPs over time, which was obtained by subtracting averaged ERP for each of the target EPRs. SNR and noise power at each electrode were averaged across subjects, respectively. Black dots indicate the position of EEG electrodes.

### 5.4 Statistical analysis

We employed two-sided Wilcoxon signed-rank test to conduct a statistical analysis on a particular dependent variable (DV) with decoding model as an independent variable (IV) for each day. A DV here was either decoding accuracy (CB:org vs. CF:org) or differences in decoding accuracy (CF:org-CB:org vs. CF:rSDL-CB:org). We applied the same test to the cosine distance of features as a DV and the ERP type (original or reconstructed) as an IV. We also conducted Friedman test for a statistical analysis of SNR (DV) with day (1, 2, 3, 4, 13+ α) as an IV for each electrode.

Prior to conducting both Friedman test and Wilcoxon signed rank test, the normality test and equality test in variance were carried out based on the Kolmogorov-Smirnov and Bartlett’s tests. It was determined that normality or equality in variance was violated for all statistical analyses. Multiple comparisons were adjusted using the Holm-Bonferroni correction.

## 6. Results

### 6.1 Decreased performance by calibration-free decoding

We compared the decoding accuracy between CF:org and CB:org (Figure 4). On three days out of four, we observed decreases in decoding accuracy when using CF:org. On day 13+α, we observed a significant decrease in accuracy using CF:org (86.67% [71.19, 91.26] MEDIAN % [99% confidence interval (CI)]) compared to using CB:org (93.33% [87.15, 94.95]) (p<0.05). Also, we observed a marginally significant decrease in accuracy using CF:org (90.00% [84.95, 93.30]) compared to using CB:org (93.33% [89.31, 95.86]) (p = 0.0726) on day 2, or using CF:org (83.33% [77.58, 88.74]) compared to using CB:org (90.00% [85.89, 92.71]) (p = 0.0726) on day 4. These findings, detailed in Table 1, demonstrated that the performance of P300-based BCIs could deteriorate without daily calibration.

**Table 1.** Individual decoding accuracy for different decoding methods across days (%)

| Methods | subject | day1 | day2 | day3 | day4 | day13+$\alpha$ | subject | day1 | day2 | day3 | day4 | day13+$\alpha$ |
| --- | --- | --- | --- | --- | --- | --- | --- | --- | --- | --- | --- | --- |
| **Calibration-based** |  | 83.33 | 86.67 | 86.67 | 90.00 | 83.33 |  | 93.33 | 93.33 | 96.67 | 93.33 | 93.33 |
| **Calibration-free** | **sub01** | 83.33 | 70.00 | 73.33 | 90.00 | 80.00 | **sub11** | 93.33 | 96.67 | 86.67 | 86.67 | 90.00 |
| **Proposed** |  | 86.67 | 86.67 | 90.00 | 93.33 | 93.33 |  | 100.00 | 96.67 | 96.67 | 93.33 | 100.00 |
| **Calibration-based** |  | 70.00 | 100.00 | 100.00 | 86.67 | 83.33 |  | 100.00 | 100.00 | 100.00 | 96.67 | 100.00 |
| **Calibration-free** | **sub02** | 70.00 | 93.33 | 80.00 | 53.33 | 50.00 | **sub12** | 100.00 | 100.00 | 96.67 | 100.00 | 96.67 |
| **Proposed** |  | 76.67 | 90.00 | 80.00 | 56.67 | 53.33 |  | 100.00 | 100.00 | 100.00 | 100.00 | 96.67 |
| **Calibration-based** |  | 93.33 | 93.33 | 90.00 | 100.00 | 100.00 |  | 96.67 | 96.67 | 100.00 | 93.33 | 96.67 |
| **Calibration-free** | **sub03** | 93.33 | 90.00 | 86.67 | 93.33 | 20.00 | **sub13** | 96.67 | 86.67 | 100.00 | 83.33 | 100.00 |
| **Proposed** |  | 93.33 | 93.33 | 83.33 | 93.33 | 23.33 |  | 93.33 | 80.00 | 96.67 | 80.00 | 93.33 |
| **Calibration-based** |  | 93.33 | 96.67 | 96.67 | 96.67 | 96.67 |  | 96.67 | 100.00 | 83.33 | 86.67 | 83.33 |
| **Calibration-free** | **sub04** | 93.33 | 96.67 | 96.67 | 93.33 | 96.67 | **sub14** | 96.67 | 93.33 | 90.00 | 83.33 | 83.33 |
| **Proposed** |  | 96.67 | 96.67 | 96.67 | 96.67 | 96.67 |  | 93.33 | 96.67 | 86.67 | 80.00 | 80.00 |
| **Calibration-based** |  | 73.33 | 86.67 | 76.67 | 86.67 | 80.00 |  | 93.33 | 86.67 | 96.67 | 86.67 | 90.00 |
| **Calibration-free** | **sub05** | 73.33 | 93.33 | 63.33 | 80.00 | 50.00 | **sub15** | 93.33 | 90.00 | 90.00 | 76.67 | 80.00 |
| **Proposed** |  | 90.00 | 96.67 | 73.33 | 86.67 | 56.67 |  | 100.00 | 93.33 | 96.67 | 93.33 | 86.67 |
| **Calibration-based** |  | 93.33 | 90.00 | 80.00 | 86.67 | 86.67 |  | 93.33 | 93.33 | 80.00 | 86.67 | 90.00 |
| **Calibration-free** | **sub06** | 93.33 | 86.67 | 80.00 | 70.00 | 93.33 | **sub16** | 93.33 | 86.67 | 83.33 | 90.00 | 90.00 |
| **Proposed** |  | 96.67 | 93.33 | 90.00 | 76.67 | 100.00 |  | 90.00 | 96.67 | 86.67 | 93.33 | 93.33 |
| **Calibration-based** |  | 93.33 | 90.00 | 63.33 | 90.00 | 96.67 |  | 100.00 | 96.67 | 76.67 | 80.00 | 93.33 |
| **Calibration-free** | **sub07** | 93.33 | 86.67 | 66.67 | 80.00 | 90.00 | **sub17** | 100.00 | 93.33 | 90.00 | 90.00 | 86.67 |
| **Proposed** |  | 96.67 | 83.33 | 60.00 | 86.67 | 90.00 |  | 100.00 | 93.33 | 86.67 | 86.67 | 83.33 |
| **Calibration-based** |  | 96.67 | 93.33 | 90.00 | 83.33 | 96.67 |  | 96.67 | 100.00 | 83.33 | 96.67 | 96.67 |
| **Calibration-free** | **sub08** | 96.67 | 96.67 | 100.00 | 90.00 | 96.67 | **sub18** | 96.67 | 83.33 | 66.67 | 76.67 | 86.67 |
| **Proposed** |  | 93.33 | 96.67 | 93.33 | 80.00 | 100.00 |  | 96.67 | 90.00 | 70.00 | 86.67 | 93.33 |
| **Calibration-based** |  | 100.00 | 100.00 | 96.67 | 96.67 | 100.00 |  | 80.00 | 80.00 | 66.67 | 70.00 | 70.00 |
| **Calibration-free** | **sub09** | 100.00 | 100.00 | 100.00 | 100.00 | 100.00 | **sub19** | 80.00 | 80.00 | 56.67 | 73.33 | 66.67 |
| **Proposed** |  | 100.00 | 100.00 | 100.00 | 96.67 | 100.00 |  | 86.67 | 80.00 | 60.00 | 63.33 | 63.33 |
| **Calibration-based** |  | 80.00 | 76.67 | 96.67 | 90.00 | 93.33 | **Average**<br>**(STD)** | 90.88<br>(9.02) | 92.63<br>(6.90) | 87.37<br>(11.31) | 89.30<br>(7.08) | 91.05<br>(8.09) |
| **Calibration-free** | **sub10** | 80.00 | 70.00 | 80.00 | 70.00 | 86.67 |  | 90.88<br>(9.02) | 89.12<br>(8.67) | 83.51<br>(13.22) | 83.16<br>(11.57) | 81.23<br>(20.82) |
| **Proposed** |  | 83.33 | 83.33 | 80.00 | 76.67 | 90.00 |  | 92.81<br>(7.14) | 91.40<br>(7.23) | 85.44<br>(13.11) | 84.91<br>(11.78) | 85.09<br>(19.86) |

### 6.2 Decoding performance without daily calibration using the proposed model

In order to assess the effectiveness of the proposed model, we assessed differences in decoding accuracy of CF:rSDL or CF:org from that of CB:org (Figure 5). We observed that a difference in accuracy between CF:rSDL and CB:org was significantly smaller than that between CF:org and CB:org on day 13+α (p<0.05). Specifically, the accuracy difference of CF:rSDL from CB:org (i.e., CF:rSDL-CB:org) was 0.00% [-15.36, 3.43], while that of CF:org from CB:org (i.e., CF:org-CB:org) was -3.33% [-19.34, -0.31]. We also evaluated whether differences in accuracy was significantly different from 0 on each day. Two-sided Wilcoxon signed-rank test with Holm-Bonferroni correction showed that difference in accuracy between CF:org and CB:org was significantly (p<0.05) or marginally significantly (p<0.1) lower than 0 on every day (days 2, 3, 4 and 13+α). In contrast, difference in accuracy between CF:rSDL and CB:org was not significantly different from 0 (p>0.1) (Figure 5). These results suggested that using CF:rSDL led to improved performance compared to using CF:org, improving the performance of calibration-free decoding closer to CB:org.

### 6.3 Visual inspection of ERP features

We visualized individual feature vectors using the t-SNE on a 2D space (Figure 6). We presented the distributions of both original ERP feature vectors before applying rSDL and reconstructed ERP feature vectors by rSDL. Figure 6 illustrates these distributions in a representative subject (sub01) who showed the most improvement of decoding accuracy from CF:org to CF:rSDL. We could observe that the reconstructed ERP features exhibited clearer separation between target and non-target over different days than the original ERP features.

### 6.4 Evaluation of feature consistency across days

The cosine distance between the original ERP features and the ERP features reconstructed by rSDL was examined to evaluate the consistency of features (Figure 7). A two-sided Wilcoxon’s signed-rank test revealed a significant reduction in the cosine distance with the reconstructed ERP features compared to the original ERP features on each of all five days. The detailed result of the cosine distance is summarized in Table 2. This cosine distance result demonstrated the efficacy of rSDL to make features in test sessions from subsequent days closer to those obtained in the calibration session on day 1, providing a clue of how the proposed method could improve calibration-free decoding performance.

**Table 2.** The averaged cosine distance on each day.

| Feature type | day 1 | day 2 | day 3 | day 4 | day 13+$\alpha$ |
| --- | --- | --- | --- | --- | --- |
| **Original** | 0.85<br>[0.83, 0.89] | 0.89<br>[0.85, 0.91] | 0.91<br>[0.87, 0.93] | 0.91<br>[0.87, 0.93] | 0.90<br>[0.87, 0.93] |
| **Recon-structed** | 0.87<br>[0.80, 0.88] | 0.87<br>[0.82, 0.89] | 0.90<br>[0.84, 0.92] | 0.89<br>[0.84, 0.92] | 0.88<br>[0.84, 0.93] |
| **Significance** | *** | *** | ** | ** | ** |

Each value indicates MEDIAN [99% confidence interval (CI)]. Significant differences are denoted by asterisks: \*\* $p < 0.01$ and \*\*\* $p < 0.001$ . Two-sided Wilcoxon signed-rank test with Holm-Bonferroni correction.

### 6.5 Signal-to-noise ratio across days

The SNR of the target ERPs was analysed across days. A statistical analysis revealed no significant difference in SNR across all days on any electrode (p>0.05) (Figure 8). Additionally, the noise power of the ERPs alone was compared across days for all electrodes, and no significant difference was found (p>0.05, Friedman test corrected by Holm-Bonferroni correction). These results indicated that the quality of the target EPRs remained consistent across all days.

### 6.6 In-depth examination of performance of the proposed model

We further observed that the performance of CF:rSDL slightly exceeded that of CB:org on day 1 (by 1.93%), as shown in the difference of decoding accuracy illustrated in Figure 5. Since both CF:rSDL and CB:org were trained and tested using the same datasets on day 1, except for using different types of ERPs, this observation of improved decoding of CF:rSDL on day 1 could signify an effect of using better ERPs reconstructed by rSDL (e.g., less noisy ERPs). Then, there was a possibility that the enhanced quality of reconstructed ERPs could also contribute to an overall improvement in decoding performance using CF:rSDL. If so, it could be ambiguous whether improved calibration-free decoding by rSDL was due to the extraction of common ERP patterns or due to the enhancement of ERP quality *per se*.

To address this ambiguity, we matched the performance level of decoders between CF:org and CF:rSDL by adjusting the number of training blocks for CF:rSDL (with a minimum of 20 blocks). The performance level was evaluated using test data on the same day of training data. In this way, calibration-free decoding by rSDL would be less likely affected by enhanced ERP quality because both CF:org and CF:rSDL were forced to possess decoders with a similar quality level. Then, we conducted cross-day decoding to assess the performance of CF:org and CF:rSDL trained on day *i* applied to test data on day *j*, ∀(*i, j)* ∈(1, 2,3, 4, 13+α) (Figure 9(A)). We confirmed that decoding accuracy was not different between CF:org and CF:rSDL when *i* = *j* on each day (p>0.05, two-sided Wilcoxon signed rank test with Holm-Bonferroni correction, see diagonal entries in Figure 9(A)). Next, we compared decoding performance when *i* ≠ *j* and found that the performance of CF:rSDL was superior to that of CF:org (p<0.01) (see off-diagonal entries in Figure 9(A) as well as Figure 9(B)). This suggests that the performance improvement in CF:rSDL compared to CF:org across days was more likely attributable to successful extraction of common ERP patterns by rSDL.

**Figure 9.**
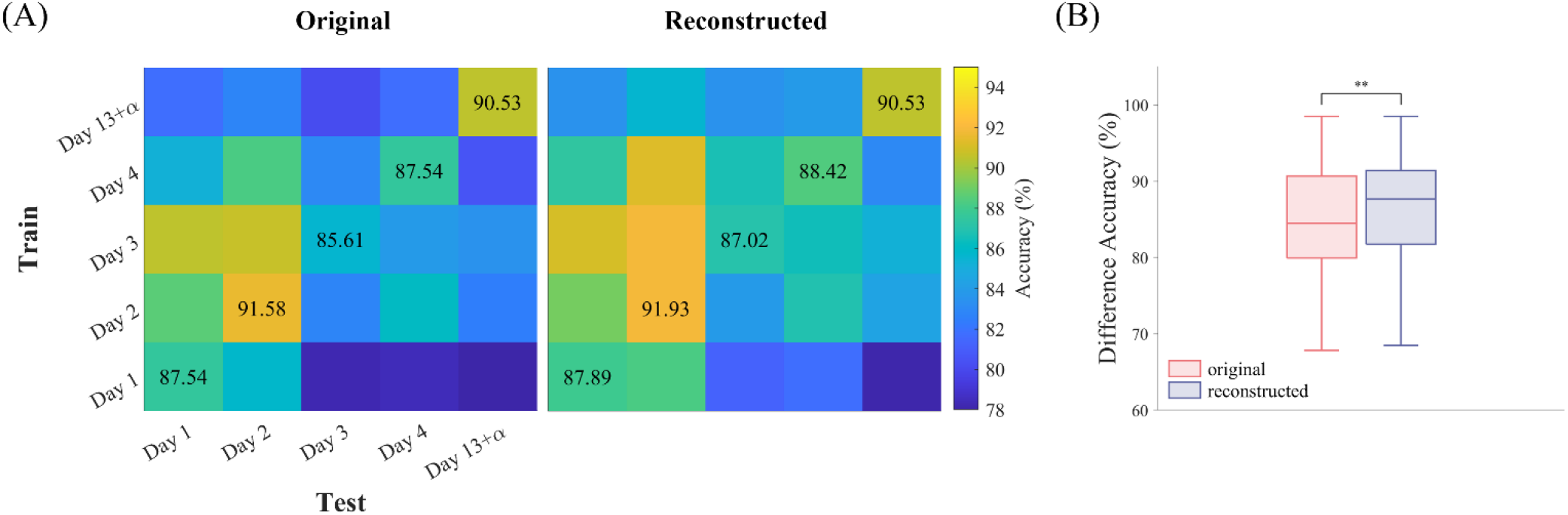
(**A**) The cross-day decoding performance using the original or reconstructed ERP features. A decoder was trained using the calibration session data on day *i* and tested using the test session data on day *j*, from which decoding accuracy was represented in the *i*-th row and *j*-th column of the matrix. The number of blocks used to train the decoder on each day was adjusted when using the reconstructed ERPs so that the diagonal accuracies became similar between the cases of using the original and the reconstructed ERPs (see the text for details). The colorbar denotes the average accuracy over subjects (%). (**B**) The box plots of cross-day decoding accuracy excluding the diagonal accuracies (i.e., same-day decoding). The cross-day decoding accuracy values in off-diagonal entries of the matrices in (**A**) were averaged for each subject. The box plots illustrate the median (horizontal line inside the box), as well as the 25th and 75th percentiles (range of the box) with a range of ± 1.5 times the interquartile range (whiskers) of the distributions of cross-day decoding accuracy over subjects. Significant differences are denoted by asterisks (** p<0.01, two-sided Wilcoxon signed-rank test with Holm-Bonferroni correction).

## 7. Discussion

In this study, we confirmed that the performance of P300-based BCIs declined over multiple days without daily calibration of decoders, indicating the necessity of daily calibration to maintain performance. To circumvent this time-consuming daily calibration, we proposed the decoding approach based on the rSDL algorithm that enabled us to infer common ERP signals. Using the reconstructed ERPs from these common ERP signals, we could improve the P300-based BCI performance without daily calibration comparable to the performance level with daily calibration. Our results demonstrate a possibility to develop a P300-based BCI without cumbersome daily calibration once it is built using the training data on the first day.

### 7.1 Did ERPs change over days?

In this study, we conducted a comparison between the performance of BCIs using the calibration-free (CF:org) and calibration-based (CB:org) decoders with the original ERPs across multiple days. The result demonstrated that the CF:org model exhibited inferior performance compared to the CB:org model. It can be attributed to the potential inconsistency of brain signals across different days. Chen *et al* showed that EEG signals could exhibit instability over time which was related to the variability of cognitive functions, such as working memory and attention, over a period of one month [59]. Moreover, as our BCI system mainly included the P300 component, it is important to consider the characteristics of this component. According to the contextual updating theory, the evaluation of task-related stimuli elicits the P300 component, which can be influenced by attentional resources and working memory [60, 61]. The P300 component can vary temporarily even over repeated blocks as attentional states change [25, 26]. As such, personal attentional states would be likely to change over multiple days. These alterations in attentional states might lead to the daily variability of the P300 component. However, further investigations are needed to gain a better understanding of the temporal variability of the P300 component in relation to attentional states as well as other cognitive states over days and even over months.

EEG signals in general exhibit variation due to various noise factors, including impedance, electrode disposition, and environmental conditions. These noise factors have the potential to subsequently affect the performance of P300-based BCIs over days. We investigated this possibility by examining the SNR across days. The result showed that the SNR and noise level did not exhibit significant variation across days. It suggests that signal quality was not a major factor contributing to the BCI performance changes across days observed in our study.

### 7.2 Was our solution effective and why did it work?

We assessed the degree of similarity in target ERPs between the calibration session on the first day and the test session on each day using cosine distance. The reconstructed ERPs by rSDL were closer between the calibration and test sessions on each day than the original ERPs (see Figure 7). The result suggests that the proposed model may provide ERPs that are relatively more consistent from training to testing across different days. This consistency is achieved by finding the atoms that represent basis ERP waveforms in EEG signals on the first day. Therefore, utilizing sparse coefficients for dictionary to determine solely whether the atoms are present in the original ERPs acquired on subsequent days and reconstructing ERPs by combining those atoms could eliminate possible variations of ERPs uniquely observed on a given day. Thus, our approach is able to reconstruct ERPs on different days by effectively utilizing the basis waveforms present on the first day.

The enhanced BCI performance over days using the proposed model may be attributed to the improvement of decoder training by using the reconstructed ERPs potentially with less noise. We investigated this confounding factor by rebuilding the decoder using reconstructed ERPs with a less amount of training data so that its decoding performance was on par with the decoder using original ERPs. Despite the comparable decoding performance in the test session on the same day as the calibration session between these two decoders, the decoder with reconstructed ERPs showed superior performance to that with original ERPs across days (see Figure 9). It suggests that our approach primarily facilitated the improvement of feature consistency across days, rather than enhancing decoder training.

In addition, in the proposed method, we employed an under-complete dictionary. The use of the under-complete dictionary has been shown to be beneficial in preventing the extraction of noise templates in a common dictionary and improving classification performances [39, 40]. Therefore, we proposed using the under-complete dictionary of ERP waveforms to address the daily calibration issue.

### 7.3 Advantages of using our approach to address daily calibration issue

To the best of our knowledge, a few research related to removing calibration for the time axis was considered regarding to P300-based BCI. Barachant and Gongedo considered the calibration problem by offering a generalized source in cross-day [28]. They obtained sessions on 6 different days, but to train variability over days to the decoder an approach of increasing the number of days for calibrating was used. Those approaches can be incompletely addressed the challenge of time-consuming daily calibration. To apply those approach in the BCI, it is necessary to obtain short signals for different days before controlling the home appliance. However, we introduced method that operate effectively without the necessity of short signal acquisition on different days. Once the dictionary was learned on the first day, it could generate more consistently for new single ERP and could use BCI immediately. Our approach offers a more time-efficient than the other approach of acquiring data over several days to remove the inherent daily fluctuations.

### 7.4 Long-term stability of P300-Based BCI

Prior research has indicated that P300-based BCI can be effectively used over extended periods by both patients and healthy individuals, necessitating repeated calibration [7, 27, 62]. Indeed, prior research has indicated that the performance of P300-based BCIs gradually increased with repeated usage [27]. It could be associated with increased user familiarity. However, this study recalibrated a decoder every day. We also observed that the P300-based BCI system with daily calibration (CB:org) maintained stable performance over days in subjects with minimal BCI experiences, demonstrating that recalibration could sustain a consistent level of BCI efficacy across days.

Our results suggest that the initial ERPs observed on the first day were present to some extent up to 15 days later, which seems to enable the continued use of BCI without daily recalibration despite the inherent variability in ERPs. However, we also found that using BCIs without daily calibration resulted in performance degradation as much as 30% compared to performance with daily calibration. Therefore, if a P300-based BCI was used without daily calibration, addressing the instability of ERPs over days seems to be crucial for the stability of BCIs.

To the best of our knowledge, there is little empirical data showing the long-term performance of P300-based BCIs without calibration. Schettini *et al.* examined the performance of P300-based BCIs when classifying data from a session with a decoder trained on data from a different session (i.e., inter-session calibration) on the same day [63]. Their findings revealed a decrease in performance with such inter-session calibration compared to using a decoder trained on data from the same session. Our observations align with these findings, as we similarly observed a decrease in performance when utilizing the calibration-free model over successive days. However, our proposed approach promises to overcome such decreases in performance without daily calibration, thus offering a new possibility of maintaining the long-term P300-based BCIs without calibration.

### 7.5 Individual differences

We observed individual differences in the instability of the BCI performance across days. This finding is consistent with a study by Nishimoto *et al.*, which examined intra-variability in EEG waveforms during a task performed on two different days, and found that some individuals were more affected by intra-individual variability on different days [20]. Individual differences in our study might be related to differences in attentional states during the task. Prior studies have indicated that the performance of P300-based BCIs can be influenced by an individual’s attention level [64, 65]. Thus, it might be possible that individuals who exhibited more instability in their performance changed their attentional level more frequently. Further studies would be necessary to elucidate individual differences in performance related to daily variation of attention levels.

### 7.6 What’s next?

This study involved BCI operation over five days, but it would be essential to conduct studies over an extended period (e.g., more than 30 days) to thoroughly evaluate the long-term use of the P300-based BCI system. Additionally, we conducted dictionary learning encompassing all EEG electrodes. However, a dictionary learned from all electrodes may potentially remove valuable signals that are unique to specific electrode or overlook inherent differences in ERP patterns across electrodes. Incorporating such electrode-specific ERP patterns into dictionary learning warrants further exploration in follow-up studies.

Although the proposed method based on the SDL method was evaluated offline in this study, it is imperative to verify its effectiveness in online P300-based BCIs. It would be feasible to employ the proposed solution in an online P300-based BCI system, given that learning the coefficients for the dictionary to reconstruct ERPs for each block requires maximum 0.069s in our case. Also, it is noteworthy that the number of features (*N*_*f*_) after removing bad EEG channels varies over days in this study. However, it can be overcome by using an interpolation process during EEG preprocessing [42]. Evaluation of the proposed method online may consider this preprocessing step.

### 7.7 Contribution of this study

The daily use of the P300-based BCI system often necessitates the time-consuming recalibration of decoding models on each day. Addressing these concerns, our study proposes a computational method to make the P300-based BCI system free from daily calibration. The effective performance of the proposed method demonstrated by the results of this study may offer a novel solution to avoid daily calibration, contributing to the translation of the P300-based BCI system into daily real-life applications.

## Acknowledgements

This research was supported by the Challengeable Future Defense Technology Research and Development Program through the Agency For Defense Development (ADD) funded by the Defense Acquisition Program Administration (DAPA) in 2022 (No.915061201).

## Ethical statement

All subjects gave informed consent for this study, which was approved by the Ulsan National Institute of Science and Technology, Institutional Review Board (UNISTIRB-21-22-A).

